# Restoration and resilience to sea level rise of a salt marsh affected by dieback events in Charleston, SC

**DOI:** 10.1101/2022.04.18.488673

**Authors:** JL Rolando, M Hodges, KD Garcia, G Krueger, N Williams, J Carr, J Robinson, A George, J Morris, JE Kostka

## Abstract

The frequency of salt marsh dieback events has increased over the last 25 years with unknown consequences to the resilience of the ecosystem to accelerated sea level rise (SLR). Salt marsh ecosystems impacted by sudden vegetation dieback events were previously thought to recover naturally within a few months to years. In this study, we provide evidence that approximately 14% of total marsh area has not revegetated 10-years after a dieback event in Charleston, SC. Dieback onset coincided with a severe drought in 2012, and a second dieback event occurred in 2016 after a historic flood influenced by Hurricane Joaquin in October of 2015, with unvegetated zones reaching nearly 30% of total marsh area in 2017. Most affected areas were associated with lower elevation zones in the interior of the marsh (midmarsh). During the 2013 dieback event, we estimate that unvegetated midmarsh area expanded by 300%. Grass planting was shown to be an effective restoration practice, with restored plants having greater aboveground biomass than relict sites after two years of transplanting. A positive restoration outcome indicated that the stressors that caused the initial dieback are no longer present. Despite that, many dieback areas have not recovered naturally even as they are located within the typical elevation range of a healthy vegetated marsh. A mechanistic modelling approach was used to assess the effects of vegetation dieback on salt marsh resilience to SLR. Predictions indicate that a highly productive restored marsh (2000 g m^-2^ y^-1^) would persist at a moderate SLR rate of 60 cm 100 y^-1^, whereas a non-restored mudflat would lose all of its elevation capital after 100 years. Thus, rapid restoration of marsh dieback is critical to avoid further degradation. Also, failure to incorporate the increasing frequency and intensity of extreme climatic events which trigger irreversible marsh diebacks underestimates salt marsh vulnerability to climate change. At an elevated SLR rate of 122 cm 100 y^-1^, most likely an extreme climate change scenario, even highly productive ecosystems augmented by sediment placement would not keep pace with SLR. Thus, climate change mitigation actions are also urgently needed to preserve present-day marsh ecosystems.

## 1. Introduction

The resilience of salt marsh ecosystems to climate change has been a source of intense scientific debate, often focused on the response of marshes to accelerated sea level rise (SLR) (Donnelly and Bertness, 2001; Kirwan et al., 2010; Kirwan et al., 2016a; Kirwan and Megonigal, 2013; Morris et al., 2002; Langston et al., 2021; Raposa et al., 2017; Rogers et al., 2019; Schuerch et al., 2018; Weston, 2014). Ultimately, the capacity of present-day marshes to persist under accelerated SLR will depend on the ability to increase in surface elevation at a rate that at least matches that of SLR (Morris et al., 2002; Reed, 1995). Salt marsh surface elevation is determined by the balance between sediment deposition and erosion, organic matter sequestration, and land subsidence (Kirwan and Megonigal, 2013; Morris et al., 2016; Mudd et al., 2009). Plant productivity is a key variable modulating marsh vertical accretion not only by accruing organic matter from belowground biomass, but also by facilitating the deposition of suspended matter during tidal inundation (Baustian et al., 2012; Mudd et al., 2009, 2010). Moreover, relative marsh elevation or elevation capital (Cahoon and Guntenspergen, 2010), the distance of the wetland surface to the lowest elevation at which plants can survive, largely determines the vulnerability to SLR (Morris et al., 2021).

In the last 25 years, sudden salt marsh dieback events in which extensive areas of vegetation die-off in short periods of time have been increasing in frequency with little understanding of the implications for the long-term resilience of marshes to SLR (Alber et al., 2008; McKee et al., 2004). Sudden salt marsh dieback events have been associated with drought, and suggested to be caused by the interaction of multiple physical, chemical, and biological stressors in the Southeast and Gulf of Mexico US (Alber et al., 2008). Specifically, elevated porewater salinity, acidification of air-exposed sediment, fungal infections, and increased snail herbivory have all been reported in previous events (Alber et al., 2008; Hughes et al., 2012; McKee et al., 2004; Silliman et al., 2005). In many cases, areas affected by sudden salt marsh dieback have naturally recovered within a few months to years (Alber et al., 2008; Lindstedt et al., 2006; Linthurst and Seneca, 1980; Marsh et al., 2016). However, there are also reports of marshes that have not recovered to date (Alber et al., 2008; Baustian et al., 2012; Marsh et al., 2016), with most of these cases representing low elevation zones that experienced an irreversible regime shift to ponds, requiring active restoration for recovery (Baustian et al., 2012; McKee et al., 2004; Ogburn and Alber, 2006; Schepers et al., 2020). In contrast, little is known about the causes and fate of dieback events causing long-term vegetation loss in areas that transitioned from salt marsh to mudflat, as well as their vulnerability to accelerated SLR. Furthermore, to the best of our knowledge there are limited studies characterizing the trajectory and recovery of salt marsh dieback events at decadal timescales (e.g., Marsh et al., 2016).

At a global scale, the area of marshland that is projected to drown as a consequence of accelerated SLR has been suggested to be offset by salt marsh migration onto higher land (Kirwan et al., 2016a). Nevertheless, even if that holds true, urban salt marshes are highly vulnerable to accelerated SLR because built infrastructure associated with coastal cities acts as a physical barrier preventing marsh migration (Kirwan and Gedan, 2019; Schuerch et al., 2018). If no actions are taken to conserve present-day urban salt marshes, the ecosystem services they provide will be lost. For example, salt marshes provide a natural defense to storm surge for coastal populations and cities (Barbier et al., 2011; Temmerman et al., 2013). Up to 41% of the world’s population lives on the coastline, with 630 million people living below projected flood levels for 2100, and lower income populations from underrepresented minority communities are more likely to live in flood prone zones (Kulp and Strauss, 2019; Martínez et al., 2006; Chakraborty et al., 2014; Eisenman et al., 2007). Conservation of urban marshes represents an effective climate change adaptation strategy with increasing relevance over time, as the frequency and intensity of tropical storms and hurricanes are already increasing (Brown et al., 2019; Paerl et al., 2019). To the best of our knowledge, little information is available on the vulnerability or resilience of urban salt marshes to accelerated SLR. Thus, there is great need to investigate the potential response of urban marshes to various climate change, ecological, and management scenarios in order to support adaptive management of the coastline.

This study was initiated to assess the vulnerability and restoration potential of salt marshes in Charleston, SC, motivated by members of the Ashleyville Neighborhood Association in West Ashley, a densely populated low-lying area that borders on urban marshes. Salt marshes in the area serve as essential habitat for many fish and shellfish species (e.g., red drum, sharks, oysters), and the National Fish and Wildlife Foundation (NFWF)-funded Coastal Resilience Assessment prioritized the West Ashley area for restoration because its habitats are generally in relatively poor condition, wildlife is exposed to high stress, and human assets show a very high vulnerability to storms along with SLR (Crist et al. 2019). Thus, we used this case study to expand the current knowledge on the trajectory of salt marsh dieback events and their effects on salt marsh resilience to climate change-induced accelerated rate of SLR. Objectives of this study were to: i. characterize the onset, dynamic, and trajectory of salt marsh dieback events in Charleston, SC ii. correlate the onset of dieback with climatological and salt marsh geophysical properties, iii. evaluate the effectiveness of salt marsh restoration by grass planting in an unvegetated marsh affected by a dieback event, and finally iv. model the resilience of dieback affected salt marshes to SLR under contrasting restoration scenarios.

## 2. Materials and methods

### 2.1 Study Area

The study was conducted in *Spartina alterniflora*-dominated salt marshes located in West Ashley, Charleston, SC. The watershed west of the Ashley River has great cultural and historical relevance since it contains Maryville and Charles Towne Landing. Maryville and Charles Towne Landing are the sites of a Gullah Geechee community established after the American Civil War and first permanent English settlement in the Carolinas in 1670, respectively. The dieback area was first discovered by members of the local Gullah Geechee community in a salt marsh adjacent to Maryville (Fig. S1).

In order to relate the expansion of dieback areas to drought and extreme precipitation, we retrieved monthly Palmer drought stress index (PDSI) values from the South Carolina Southern Division (January 2009 to August 2021) using NOAÁs National Centers for Environmental Information database (https://www.ncdc.noaa.gov/temp-and-precip/drought/historical-palmers/), and monthly precipitation values from the Charleston area, SC (ThreadEx) using NOAA’s Online Weather Data (NOWData, https://www.weather.gov/wrh/Climate?wfo=chs). The PDSI is a widely used drought index based on the supply and demand of a water balance equation of precipitation, evapotranspiration, soil moisture loss and recharge, and runoff, which has been used to study the effects of water limitation on agricultural and ecological systems (Palmer, 1965).

### 2.2 Characterization of marsh dieback trajectories using multispectral high-resolution imagery

Multispectral imagery was acquired from three sources, satellite imagery from the PlanetScope constellation (Planet Team, 2017), and MAXAR’s WorldView-3 satellite, and aerial imagery from US Department of Agriculture’s (USDA) National Agriculture Imagery Program (NAIP). The NAIP dataset consists of 4-band [blue, green, red, and near infrared (NIR)], 1-meter resolution imagery (http://www.fsa.usda.gov/programs-and-services/aerial-photography/imagery-programs/naip-imagery). The PlanetScope constellation acquires daily, 3-meter resolution imagery, using a 4-band (blue, green, red, NIR) sensor. We used the Planet’s Surface Reflectance product, processed to top of atmosphere reflectance, and atmospherically corrected to surface reflectance (Planet Team, 2019). PlanetScope images were acquired during low tide, and on cloudless days. The MAXAR WorldView-3 acquires 4-band (blue, green, red, NIR), 30-cm resolution images. The used View-Ready WorldView-3 image was retrieved during low tide on a cloudless day, was radiometrically and sensor corrected, and atmospherically corrected to surface reflectance (MAXAR, 2021). The complete imagery used in this study consisted of 9 images retrieved from 2009 to 2021 (further details on Table S1).

Two vegetation indices were calculated from all acquired images: the normalized difference vegetation index (NDVI), and the modified soil-adjusted vegetation index 2 (MSAVI2):

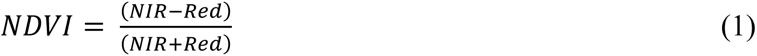

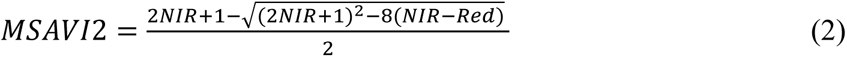

Total unvegetated area was estimated for all NAIP images, and the 2016 WorldView-3, and 2021 PlanetScope satellite images. We visually inspected the coordinates of vegetated and unvegetated points (median observations per year: 218) using high-resolution images at a scale of 1:500 to 1:1000 in QGIS 3.16.5 (QGIS.org, 2021). Raster values of the four bands and NDVI were extracted from the pixels containing the vegetated and unvegetated points. Random forest models were used to estimate the probability that a pixel was unvegetated using the four bands and NDVI as explanatory variables. Out-of-bag estimate of error rate was lower than 3.5% for all models. Pixels with a greater than 75% probability of being unvegetated were defined as such, while pixels with a lower than 25% likelihood of being unvegetated were defined as vegetated areas. This analysis was performed using the raster, and randomForest R packages in the R v. 4.1.0 environment (Hijmans and van Etten, 2012; Liaw and Wiener, 2002; R Core Team, 2021).

In order to investigate if unvegetated areas were preferentially associated with creek bank or midmarsh zones, we drew all creek bank edges from the study area (Fig. S1). Edges were drawn using the 2019 NAIP aerial image and the corrected 2017 LIDAR derived DEM at a scale of 1:250 to 1:500 in QGIS 3.16.5 (QGIS.org, 2021). We generated a raster measuring the distance of each pixel to the closest creek bank edge using the “Join attributes by nearest” algorithm in QGIS with a resolution of 1 m. Based on the well-known natural zonation of *S. alterniflora* (Mendelssohn and Morris, 2002), with taller plants found closer to tidal creeks, areas within 7.5 m of a tidal creek were defined as “creek bank marsh”, whereas areas > 7.5 m distal to creeks were defined as “midmarsh” (Valiela et al., 1978). The 7.5 m threshold characterizing the ecosystem state change was defined by visually inspecting an aboveground biomass vs. distance from tidal creek biplot (Fig. S2, method for aboveground biomass estimation below).

### 2.3 Field sampling

Field sampling was performed during spring and summer of 2021. Two field campaigns were performed on 27-28 May, and 20-26 July of 2021. We sampled 60, and 119 locations, in May and July, respectively (total = 179 points). Field sampling was performed non-randomly by establishing transects across gradients of *S. alterniflora* biomass from the creek bank to the interior of the marsh (Fig. S1). The dieback area adjacent to Maryville was oversampled due to our interest as a potential restoration site. At all locations (n = 179), a high-accuracy real time kinematic (RTK) survey was conducted using a Trimble GNSS GPS receiver (Trimble, Sunnyvale, CA, USA). Elevation was expressed in meters in the North American Vertical Datum of 1988 (NAVD88) GEOID 18. Instrument accuracy is of 2 cm root mean square error (RMSE), horizontal and vertical. During the July field sampling, *S. alterniflora* shoot density and height were measured using a 0.25 m^2^ quadrat at 82 points sampled in parallel for RTK surface elevation. Shoot density was measured by counting all *S. alterniflora* stems inside the 0.25 m^2^ quadrat. Shoot height was measured randomly from 6 stems per quadrat. In order to estimate aboveground biomass, 71 *S. alterniflora* shoots of known height were harvested from 37 sampling points across contrasting *S. alterniflora* growth zones. Harvested shoots were oven-dried at 60 °C for one week and their shoot height and dry weight were used to construct a biomass allometric equation. For all sampling points, biomass was calculated for individual shoots using the constructed allometric equation, and mean shoot biomass (n = 6) multiplied by shoot density to calculate aboveground biomass at the plot level (g m^-2^).

### 2.4 Acquisition of LIDAR derived DEM

A light detection and ranging (LIDAR) derived bare-earth digital elevation model (DEM) constructed from LIDAR aerial acquisition conducted from February 25, 2017 to March 09, 2017 in Charleston County was used in this study (South Carolina DNR, 2018). LIDAR acquisition involved Axis GeoSpatial (Easton, MD, USA) operating a Cessna 206H outfitted with a Riegl LMS-Q1560 dual channel laser scanner system at 2043 m above ground level. Scanner Pulse Rate was 800 kHz with unlimited number of returns per pulse. The DEM vertical datum used was NAVD88 Geoid 12B and was projected to the NAD83 (2011) South Carolina State Plane at a pixel resolution of 1 m. System parameters for LIDAR acquisition are described in Table S2 (further details can be found in South Carolina DNR, 2018).

### 2.5 Correction of the LIDAR derived DEM

High-accuracy RTK elevations were used to correct the bare-earth LIDAR derived DEM. Error of the LIDAR derived DEM was defined as: Z_LIDAR_ - Z_RTK_ for each RTK observation, where Z is NAVD88 elevation. We used a multispectral PlanetScope satellite image retrieved 5 days before LIDAR acquisition (02/20/2017) to correct the DEM based on the known relation between LIDAR error and *S. alterniflora* height (Hladik and Alber, 2012). The 179 RTK observations were randomly split for model training and testing in a 7 to 3 ratio, respectively. Error of the LIDAR derived DEM was fitted to a multiple lineal regression model using automated model selection based on Akaike Information Criterion (AIC) as implemented in the glmulti R function (method = “h”) (Calcagno and de Mazancourt, 2010). We used the four bands of the PlanetScope image (Red, Green, Blue, and NIR), two vegetation indices (NDVI, and MSAVI2), and the original LIDAR derived DEM elevations as potential explanatory variables. The DEM was corrected by solving the multiple linear regression for the RTK observed elevation. Performance of the best model was tested based on root mean square error (RMSE) calculated as: *RMSE* = 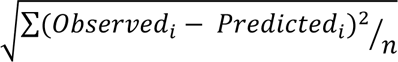 where *Observed_i_* is the ith observation of the RTK elevation, and *Predicted_i_* is the ith observation of the LIDAR derived DEM (Maune et al., 2007). We calculated the RMSE for the test and train datasets of both the corrected and uncorrected LIDAR derived DEM.

### 2.6 Estimation of 2021 aboveground biomass

Aboveground biomass was estimated for the total study area based on *in situ* biomass observations made during the July 2021 field campaign. Aboveground biomass (g m^-2^) estimated from 82 sampling points (section 2.3) were randomly split into train and test datasets (80% and 20%, respectively). A random forest model was constructed using the corrected 2017 LIDAR derived DEM, MSAVI2, NDVI, and the green, red and NIR bands from a PlanetScope image (acquired in 08/01/2021) as explanatory variables. The model grew 5000 trees; and in each split, 2 variables were randomly sampled as candidates. The performance of the model was assessed by calculating the RMSE of the train and test datasets (as in section 2.5). A raster containing the predicted aboveground biomass for the whole study area was constructed using the random forest model. Aboveground biomass was set to 0 in all unvegetated areas (as inferred in section 2.2). The model was built using the raster and randomForest R packages in the R v. 4.1.0 environment (Hijmans and van Etten, 2012; Liaw and Wiener, 2002; R Core Team, 2021).

### 2.7 Pilot grass planting restoration

A pilot restoration project led by the South Carolina Department of Natural Resources (SC-DNR) was initiated in the Maryville marsh in 2019 in order to test the effectiveness of marsh restoration in long-term affected ecosystems impacted by dieback events. The restoration effort consisted of *S. alterniflora* planting using seedlings grown from local sources. Botanical seeds were annually collected in October thru December each year (2019, 2020, and 2021) from 11 unique sites, all located in Charleston County. After collection, seeds were kept at 4°C for eight weeks, after which they were germinated in planting trays filled with fresh water substrate. After 6-7 months, seedlings ∼55.0 cm tall were transplanted into the dieback area from May to August. When transplanting, a spacing of 30.5 cm was used. Restoration was conducted in 6 individual patches with dimensions ranging from 48.6 m^2^ to 183.3 m^2^. Plots from transplants performed in 2019, 2020 and 2021 were studied. In order to assess the effectiveness of the pilot restoration project, aboveground biomass was estimated during the July field campaign in 5 and 12 quadrats (0.25- m^2^) from restored plots planted in 2020 and 2021, respectively. In order to have a complete and balanced statistical design at the end of the growing season, a second field assessment was performed in September 22^nd^ 2021. In this assessment, 10 quadrats per plot were sampled in plots planted in 2019, 2020, and 2021, and two patches of adjacent relict marsh. Aboveground biomass was calculated based on shoot height and density as previously described (section 2.3). A one-way ANOVA with the restored/natural plots as a fixed factor was used to explain differences in *S. alterniflora* aboveground biomass for both the July and September assessments. In case of finding statistical significance (p < 0.05), a Tukey’s post-hoc test using the *emmeans* R package was performed (Lenth, 2016).

### 2.8 Modeling of marsh elevation under primary production, management, and sea level rise scenarios

The Marsh Equilibrium Model (MEM) v. 9.01 (Morris et al., 2002, 2021) was employed to assess the resilience of the Charleston salt marsh to accelerated SLR under different scenarios of (i) *S. alterniflora* primary productivity, (ii) human intervention (none or thin layer placement), and (iii) rates of SLR (regional NOAA’s intermediate low of 60 cm in 100 y^-1^ and intermediate of 122 cm in 100 y^-1^). The rationale to utilize the MEM was to assess the long-term effect (100 y) of marsh vegetation loss with and without restoration in Charleston, SC. The MEM is a one-dimensional mechanistic model that projects change in marsh elevation with SLR as a function of primary production and sediment accretion. It was modified to simulate thin-layer placement (TLP) of mineral sediment by allowing the user to specify periodic sediment additions to a specified thickness, net of compaction. The model was coded in VBA behind an Excel user interface.

We simulated two rates of SLR, based on regional NOAA’s intermediate low (60 cm in 100 years), and intermediate (122 cm in 100 years) projections by 2100 (Sweet et al., 2017). The intermediate low scenario has a 73% and 96% probability of being exceeded in 2100 under RCP4.5 and RCP8.5 scenarios, respectively; while the intermediate scenario has a 3% and 17% probability of being exceeded by 2100 under RCP4.5 and RCP8.5 scenarios, respectively (Sweet et al., 2017). Starting sea-level rise rate was set as 3.4 mm y^-1^, as calculated for the Charleston Harbor by Morris and Renken (2020). The elevation range of marsh vegetation was assumed to be within 25 cm below mean sea level (MSL), and 30 cm above mean high water (MHW), as characterized in South Carolina salt marshes (Morris et al., 2013). Tidal range, MSL and MHW during the study period (Jan 2009 to August 2021) was 76 cm, 4 cm NAVD88, and 80 cm NAVD88, respectively. Optimal elevation was determined as 5 cm above MSL based on a biplot of elevation and aboveground biomass from the study area (Fig. 4f). Four different optimal values of aboveground primary productivity were modelled: 0, 500, 1000, and 2000 g m^-2^ y^-1^ in order to assess the effects of plant productivity and restoration by grass planting on marsh resilience to SLR. The median elevation of all dieback pixels in 2021 was set as the initial elevation: 0.478 m NAVD88. Total suspended matter (TSM) was estimated as 23 mg L^-1^ from remotely sensed GlobColour OLCIA imagery (http://hermes.acri.fr/) as the average of monthly values from 2016 to 2021. TSM was extracted from the closest pixels to the southern border of our study area in the Charleston Harbor. Finally, a treatment with 5 cm of thin layer placement (TLP) every 25 years was included into the study to simulate a more aggressive restoration scenario. Thin layer placement consists of spreading sediment onto the marsh platform to increase surface elevation (Ford et al., 1999). After TLP, marsh vegetation was simulated to be completely lost, and fully recovered 10 years after disturbance.

## 3. Results

### 3.1 Salt marsh dieback trajectory

We observed an expansion of large unvegetated areas in 2013 that coincided with a severe drought in 2011-2012 (Fig. 1). Charleston, SC, experienced its most severe drought over the last 100 years in 2011-2012, concurrent with an over 200 % increase of unvegetated marsh area (64 ha, Fig. 2a). The South Carolina Southern NOAA division, which contains the city of Charleston, experienced an extreme drought (PDSI < -4.0) from June of 2011 to April of 2012 (Fig. 1). A second large expansion of unvegetated area was observed in 2016-2017 which was not drought-induced. The onset of the 2016-2017 dieback event coincided with the historic flood of the city of Charleston caused by record precipitation associated with the tropical cyclone Joaquin in October 2015 (Fig. 1). In 2019, the vegetated area of West Ashley had recovered up to 2013 levels which have persisted since. Based on field inspection and aerial imagery, we found that the affected salt marsh ecosystem transitioned from vegetated salt marsh to mudflat after both dieback events.

**Figure 1:**
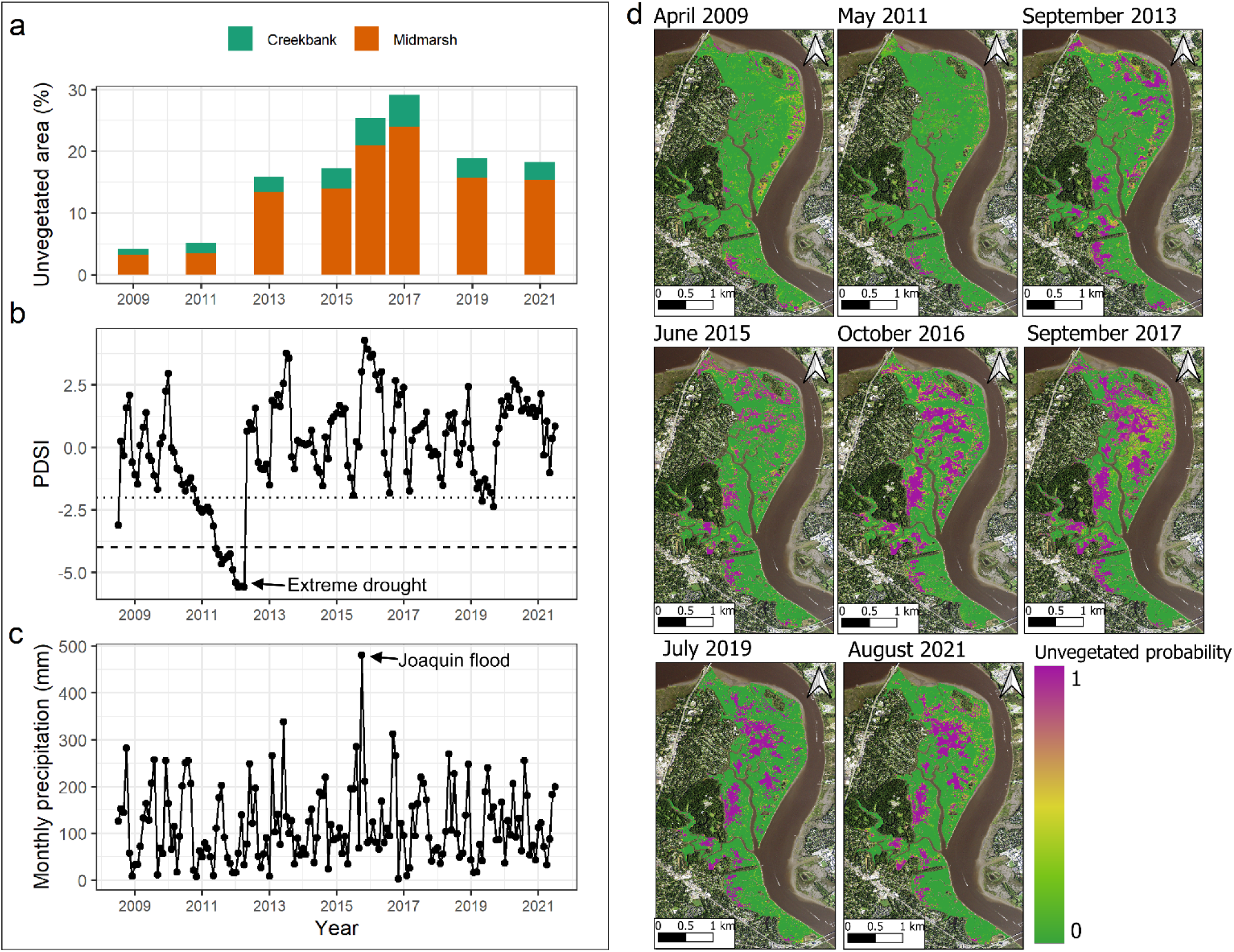
Dieback trajectory of the saltmarsh located in West Ashley, Charleston, SC. **a** Percentage of unvegetated marsh area over time. Total area is partitioned into creek bank or midmarsh categories based on areas closer or further than 7.5 m to its closest creek bank, respectively. **b** Palmer drought severity index (PDSI) from the South Carolina Southern Division the over time. Values lower than -2 represent moderate drought (dotted line), while values lower than -4 extreme drought conditions (dashed line). **c** Monthly precipitation from the Charleston, SC area retrieved from NOAA’s Online Weather Data (NOWData). **d** Maps showing the probability of an area being unvegetated. Based on individual random forest models per image.

**Figure 2:**
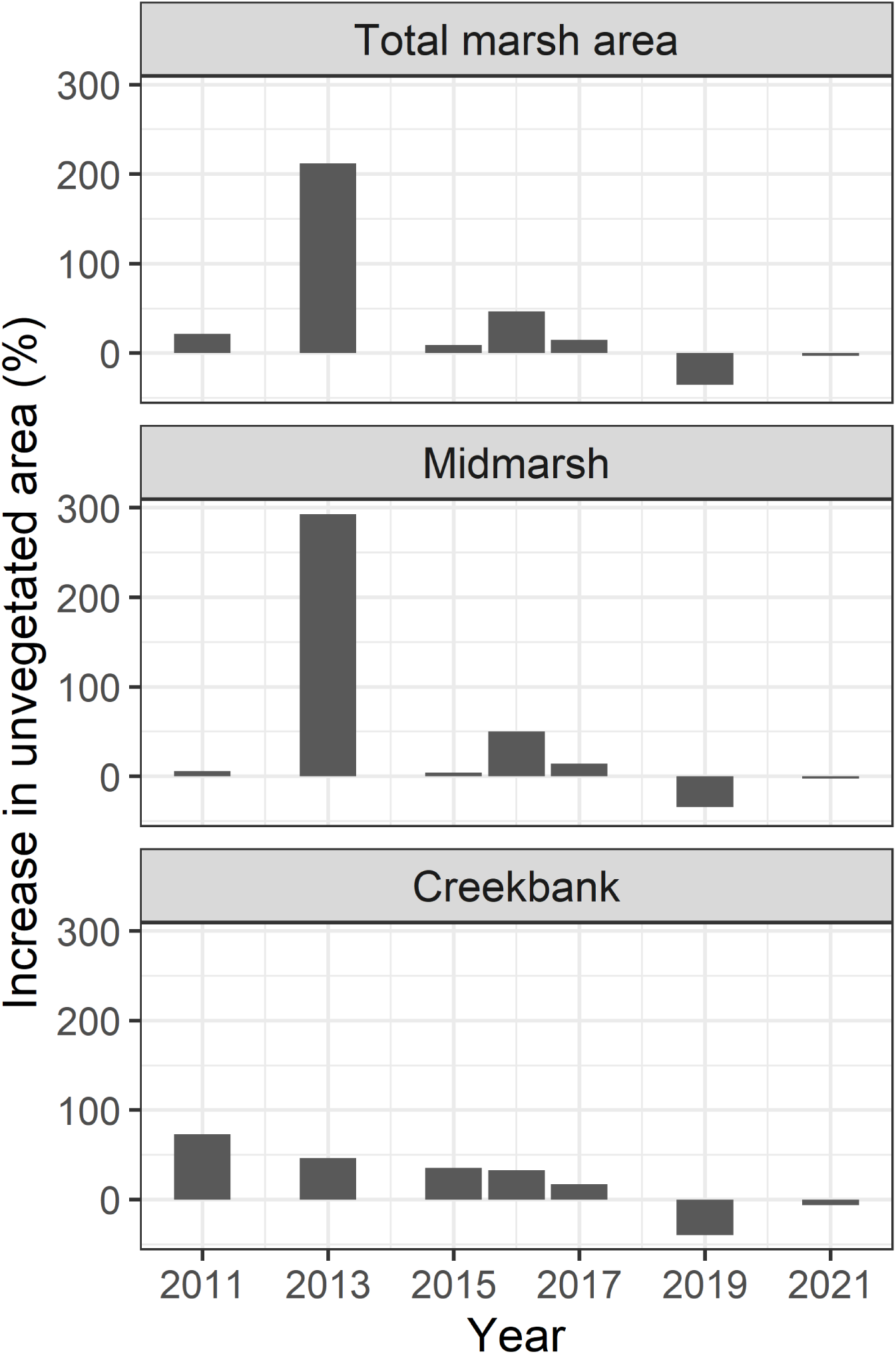
Dieback events were associated with an expansion of unvegetated area in the midmarsh zone. Barplots showing the percent increase in unvegetated areas since the last evaluation (1 or 2 years). Analysis performed for the total marsh (a), midmarsh (b), and creekbank (c) areas.

The dieback of 2013 caused a 300% expansion of unvegetated area in the interior of the marsh (Fig. 2b). In the 2016 event, the midmarsh and creekbank experienced a 50% and 33% increase of unvegetated area, respectively (Fig. 2b,c). Moreover, from 2011 to 2017 there was a constant loss of about 50% of creekbank salt marsh every two years (Fig. 2c). A LIDAR derived DEM acquired on 2017 was corrected for the studied area, improving the error estimation by approximately 30% on both training and testing datasets (RMSE after correction: less or equal to 8 cm in both train/test datasets, more detailed information in Supplementary Text S1, and Fig. S3). Using the corrected 2017 LIDAR derived DEM, we observed that both creek bank and midmarsh unvegetated areas were associated with lower elevation terrain (Fig. 3). The difference in median elevations between vegetated and unvegetated areas ranged from 13.3 cm to 27.4 cm, and 6.3 cm to 14.3 cm in creek bank and midmarsh zones across years, respectively. Since the difference in elevation could have been caused by land subsidence after the dieback event, we repeated the analysis with an uncalibrated 2009 LIDAR derived DEM (South Carolina DNR, 2009), finding similar results (Fig. S4). In the 2009 DEM, difference between median elevation from vegetated and unvegetated areas ranged from 4.4 to 23.0 cm, and 2.4 to 16.3 cm in creek bank and midmarsh zones, respectively. The results suggest that lower elevation persisted in the unvegetated areas prior to the dieback event.

**Figure 3:**
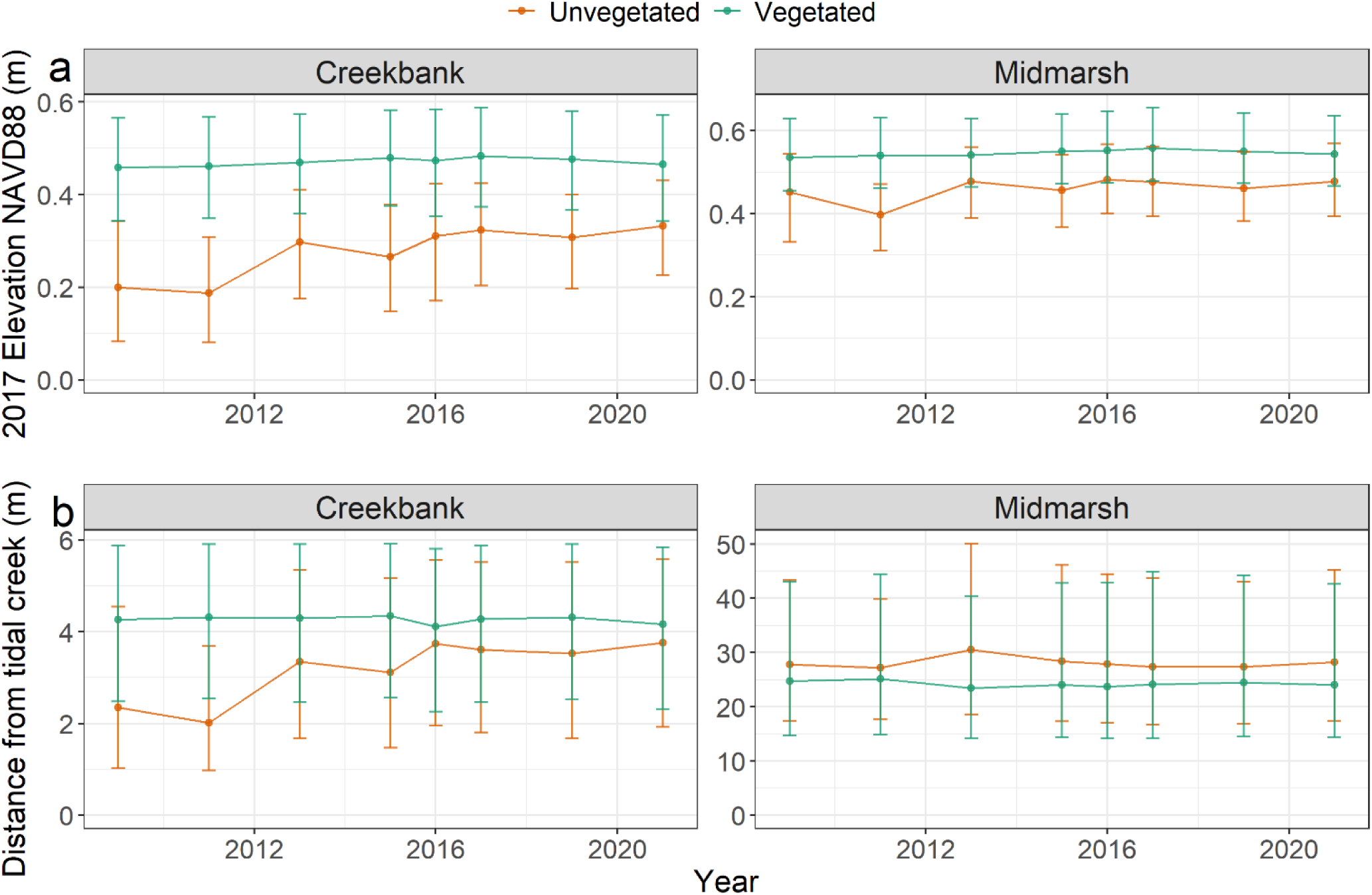
Characterization of unvegetated and vegetated areas within the West Ashley, SC salt marsh. **a** Median ± interquartile range of the field-corrected 2017 light detection and ranging (LIDAR) derived digital elevation model (DEM) of vegetated and unvegetated areas over time. **b** Median ± interquartile range of the distance from the closest tidal creek of vegetated and unvegetated areas over time.

**Figure 4:**
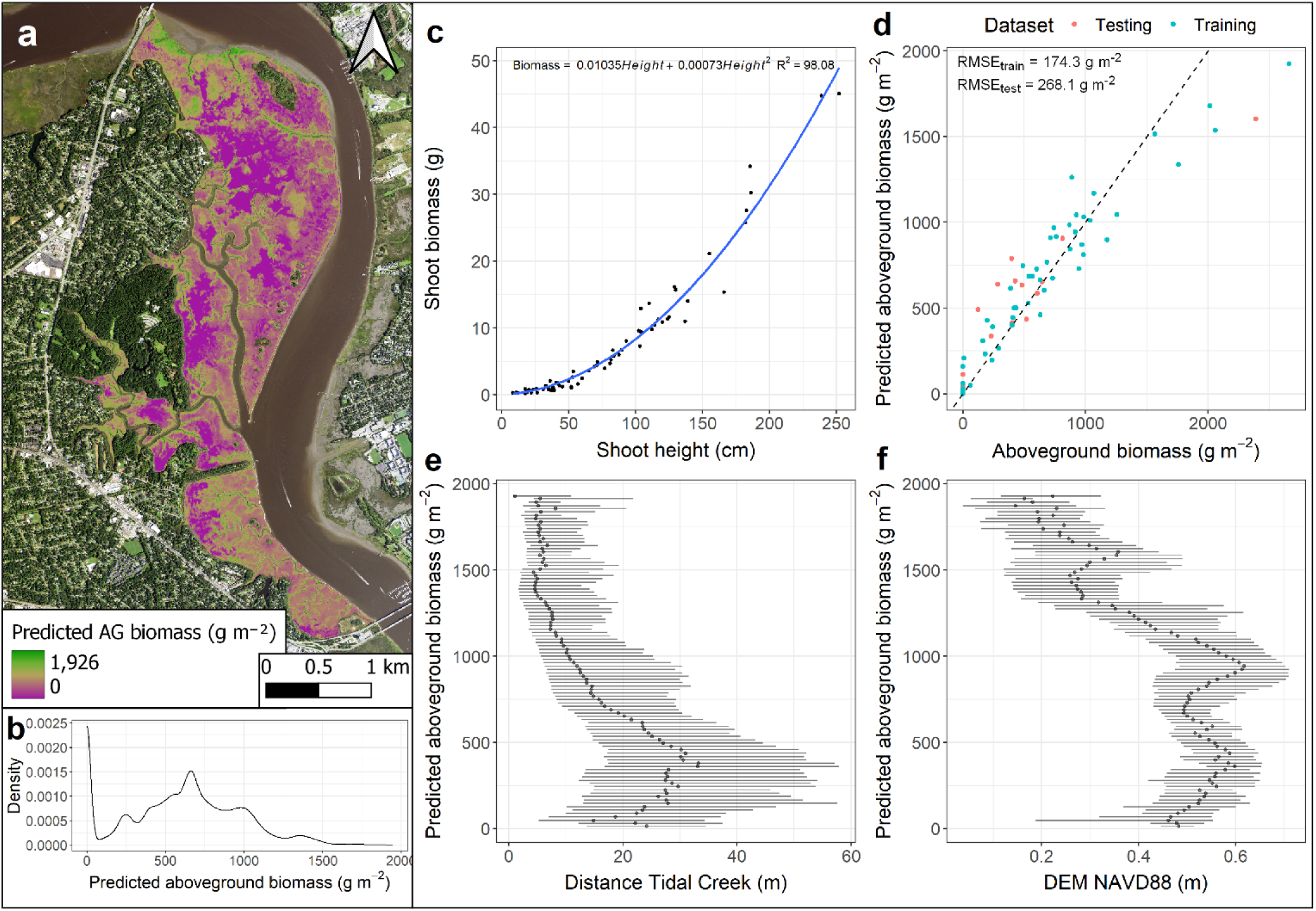
Aboveground biomass (AGB) estimation of the West Ashley, SC salt marsh during July 2021. **a** Map of AGB based on a random forest model using the 2017 elevation digital elevation model, vegetation indices and bands from a PlanetScope 4-band image. **b** Density plot of the predicted AGB values. **c** Allometric biomass equation based on shoot biomass. **d** Goodness of fit of the random forest model in the testing and training datasets. Relation between predicted AGB and distance from tidal creek **(e**), and the 2017 elevation model (**f**). Points and error bars represent AGB median and interquartile range on equidistant increments, respectively.

### 3.2 *Spartina alterniflora* primary production

A site-specific allometric equation was constructed to quantify the primary production of the Charleston salt marsh during July of 2021 (Fig. 4a,c). The random forest model used to estimate plant aboveground biomass was based on the 2017 calibrated LIDAR derived DEM, vegetation indices, and satellite bands from an August 2021 color near infrared PlanetScope image. The RMSE_test_ of the model was 268.1 g m^-2^. Median predicted aboveground biomass was 623.1 g m^-2^, ranging from 0 to 1944 g m^-2^ (Fig. 4b). Biomass was within the expected range for Charleston’s latitude (predicted end of season productivity based on Kirwan et al., 2009: 574.8 g m^-2^ y^-1^). Aboveground biomass was closely associated with elevation and distance to tidal creeks (Fig. 4e,f, Fig. S2), with greater aboveground biomass observed in lower elevation areas within closer proximity to tidal creeks (Fig. 4e,f).

### 3.3 Grass transplanting is a successful restoration method in West Ashley, SC

Guided by the South Carolina Department of Natural Resources, seedlings from botanical seeds of *S. alterniflora* were collected in Charleston, SC, propagated under greenhouse conditions, and transplanted by community volunteers in July of 2019, 2020, and 2021 in a West Ashley dieback area that has been unvegetated since 2013 (Fig. 5, S1). Aboveground biomass was determined during the July field trip for the 2020 transplanted plot compared to reference plots of natural midmarsh grass in the study area (Fig. 5). Aboveground biomass of the naturally vegetated midmarsh and the 2020 transplanted plot were non-statistically different and had an average ± S.E. of 527 ± 43 g m^-2^, and 700 ± 185 g m^-2^, respectively. Repeated measurements conducted in September showed that the restored plots planted in 2019 had greater aboveground biomass in comparison to reference relict marshes adjacent to the restoration site (Fig. 5). The plot transplanted in 2020 was non-statistically different from reference relict marshes (Fig. 5). Average ± S.E. aboveground biomass was 1726 ± 207 g m^-2^, and 1290 ± 168 g m^-2^ for the 2019 and 2020 transplanted plots, respectively.

**Figure 5.**
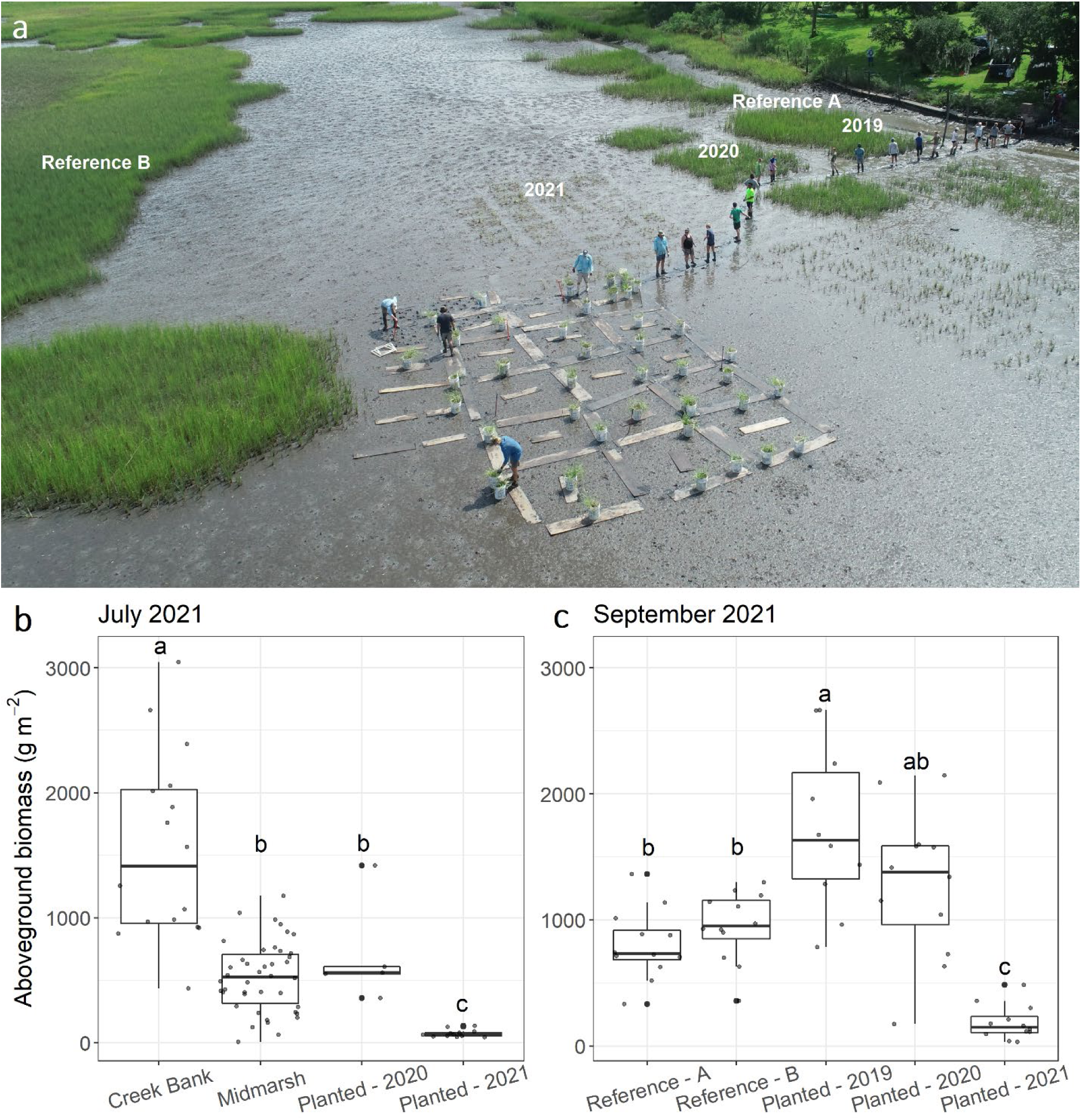
Grass planting is an effective restoration practice in Maryville, SC. **a** Drone image from the restoration site. Plots sampled are labelled on top in white. Photo acquired by the South Carolina Department of Natural Resources Shellfish Research Section on 21 July of 2021. **b** Boxplot representing the interquartile range of aboveground biomass from all sampled creek bank, midmarsh, and 2020 and 2021 restored plot during the July 2021 field campaign. **c** Boxplot representing the interquartile range of aboveground biomass from 10 quadrats sampled from the two reference sites, and the 2019, 2020, and 2021 restored plot during September of 2021. Different letters indicate statistically significant differences (p < 0.05) between treatments according to a Tukey test using *emmeans* R package (Lenth, 2016).

### 3.4 Salt marsh resilience to sea level rise under contrasting restoration scenarios

The model output indicated that mudflat areas (*S. alterniflora* primary productivity = 0 g m^-2^ y^-1^) rapidly lose elevation capital at a SLR rate of 60 cm in 100 y; decreasing in elevation by a total of 38 cm after the 100 years (Fig. 6). Restored salt marsh ecosystems with an optimal primary productivity of 500 g m^-2^ y^-1^, 1000 g m^-2^ y^-1^, and 2000 g m^-2^ y^-1^ would persist under the 60 cm in 100 y SLR scenario, but losing 37.4 cm, 33.0 cm, and 24.5 cm of elevation capital after 100 years, respectively (Fig. 6). At a SLR rate of 122 cm in 100 y, even a healthy restored salt marsh with an optimal primary productivity of 2000 g m^-2^ y^-1^ would collapse in less than 75 years (Fig. 6). Simulating an external supply of sediment (i.e., thin layer placement) at a rate of 5 cm of inorganic sediment every 25 years would maintain the elevation capital of all simulated salt marsh ecosystems in the 60 cm in 100 y SLR scenario. However, thin layer placement of a highly productive salt marsh (2000 g m^-2^ y^-1^) under the extreme 122 cm in 100 y SLR scenario would result in the persistence of the ecosystem for only 100 years (Fig. 6).

**Figure 6.**
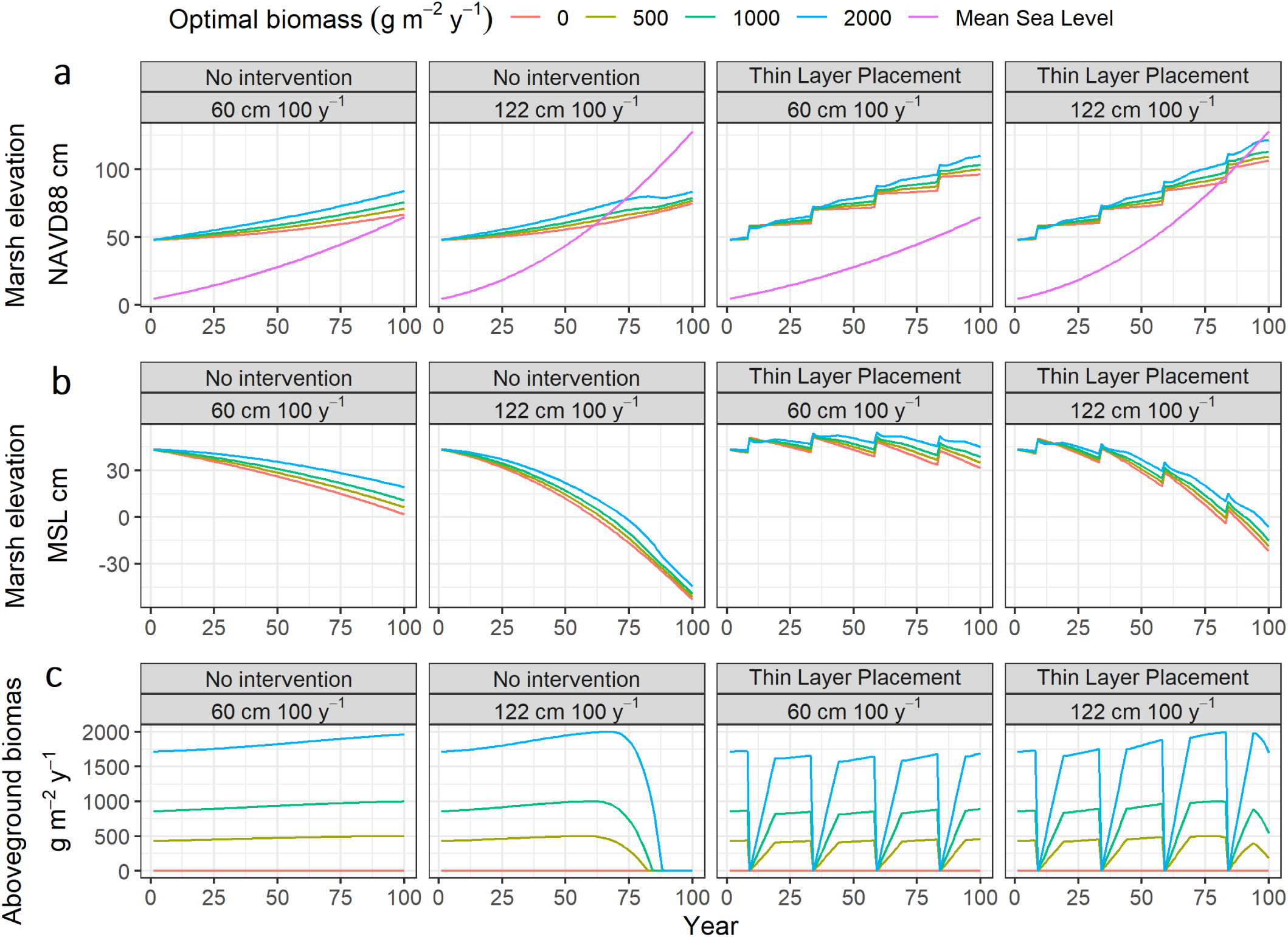
Modelled salt marsh resilience under different scenarios of sea level rise, human intervention, and plant primary production using the Marsh Equilibrium Model (MEM) (Morris et al., 2002). Modelled optimal biomass values ranged from 0 g m^-2^ y^-1^ (mudflat) to highly productive salt marshes of 2000 g m^-2^ y^-1^. Thin Layer Placement was assumed to be performed every 25 years by applying 5 cm of inorganic sediment. Vegetation was assumed to fully recover after 10 years of disturbance. The two sea level rise scenarios modelled were based on regional NOAÁs intermediate low, and intermediate of 60 cm and 122 cm in 100 y, respectively (Sweet et al., 2017). Marsh elevation (cm) over time projected to NAVD88 (**a**), and mean sea level (MSL) (**b**). **c** Modelled salt marsh aboveground biomass over time.

## 4. Discussion

### 4.1 The resilience of salt marshes to accelerated sea level rise and extreme climatic events

Extreme drought events have been linked to sudden vegetation diebacks in several salt marshes in the USA Southeast and Gulf of Mexico over the last 25 years (Alber et al., 2008; Lindstedt et al., 2006; McKee et al., 2004). The drought-induced mechanism of sudden marsh dieback has been associated with the interaction and cascading effects of multiple stressors, such as extreme porewater salinity, sediment acidification due to the oxidation of air-exposed sulfur minerals into sulfuric acid, infection by fungal pathogens, and an exponential increase of periwinkle snail herbivory (Alber et al., 2008; Hughes et al., 2012; McKee et al., 2004; Silliman et al., 2005). More recently, extreme precipitation from catastrophic tropical cyclones was suggested to trigger marsh vegetation diebacks by either intrusion of salt water into brackish/freshwater marshes and/ or by extending periods of tidal inundation which in turn exceed the physiological tolerance of marsh plants (Marsh et al., 2016; Ramsey et al., 2012; Stagg et al., 2021). The co-occurrence of biannual vegetation loss with extreme drought and flooding events in our study provides substantial evidence that marsh vegetation dieback is attributed to extreme climatic events in the USA Southeast.

Salt marsh dieback events are often described as transient phenomena (Alber et al., 2008). In contrast, we show that marsh dieback triggered by extreme climatic events can be long-lasting, with an approximate 14% loss of total vegetated area after 10 years. To the best of our knowledge, this is the first time a decadal timeframe is used to characterize a long-term salt marsh dieback event induced by an extreme climatic event. It is intriguing that the surface elevation of the dieback areas, or elevation capital, remains within the range of a hypothetically healthy marsh (Morris et al., 2002). Furthermore, we provide evidence that the marsh may be restored by grass planting, suggesting that the stressors that caused dieback may be overcome, despite the fact that the majority of dieback areas have not revegetated naturally. Potential explanations for slow or no recovery require further research and may be associated with limited seed dispersion and viability, poor vegetative reproduction, as well as disturbed marsh hydrology and geomorphology (i.e., tidal creek erosion and sedimentation limiting previously normal tidal flow).

The increasing frequency and intensity of severe climatic events due to climate change is likely to further develop marsh diebacks similar to those we observed in Charleston (Brown et al., 2019; Dai, 2013; Paerl et al., 2019). Long-term vegetation loss, as shown in this study and previous work, increases the vulnerability of salt marsh ecosystems to SLR (Baustian et al., 2012; Mudd et al., 2009; 2010; Fig. 6). Thus, the effects of extreme climatic events on marsh vegetation must be incorporated into assessments of the resilience of the ecosystem to climate change. Current models designed to project marsh resilience to accelerated SLR often assume that the ecosystem collapses due to the progressive loss of elevation relative to mean sea level (Morris et al, 2002; Kirwan et al., 2010, 2016b). In other words, the marsh drowns with the loss of elevation capital. Failing to incorporate the increasing frequency and intensity of extreme climatic events which trigger irreversible marsh diebacks will lead to an underestimation of the vulnerability of salt marshes to climate change. For example, our modelling approach forecasted that after 100 years under a moderate climate change scenario, the elevation of a highly productive marsh would exceed that of an unvegetated dieback marsh by 17.5 cm (see below for further discussion).

### 4.2 Call for restoration of urban marshes

Urban salt marsh ecosystems are more vulnerable than non-urban marshes due to their limited migration space (Schuerch et al., 2018). At the same time, the value of the ecosystem services they provide are very high due to their close proximity to heavily populated areas. Thus, their conservation and restoration have been proposed as an effective adaptation strategy to climate change, with special emphasis on their ability to attenuate storm surge and coastal erosion (Temmerman et al., 2013; Geedicke et al., 2018). Hybrid solutions in which engineered structures are combined with coastal wetlands to protect built infrastructure from storm surge were shown to be more effective than engineered defenses alone (Temmerman et al., 2013; Zhu et al., 2020). However, despite the enhanced value and vulnerability of urban salt marshes, these ecosystems remain understudied in comparison to their non-urban counterparts (Alldred et al., 2020).

Our results corroborate evidence from non-urban marshes to show that marsh grass planting is an effective restoration practice in an urban marsh that experienced dieback (Baustian et al., 2012; Linthurst and Seneca, 1980; Ogburn and Alber, 2006). The multiple stressors that lead to sudden vegetation dieback have yet to be completely revealed (Evans et al., 2021). The ability of unvegetated marshes to respond positively to grass planting is likely dependent on the strength of the marsh platform and whether it is located within the elevation range of a stable marsh domain (*sensu* Morris et al., 2002), along with sufficient sediment supply from the surrounding estuary (Baustian et al., 2012; Liu et al., 2021; Webb et al., 1995; Wilsey et al., 1992). As shown in our modelling approach and by previous work, plant primary production is a key variable modulating marsh accretion (Morris et al., 2002; Mudd et al., 2009, 2010; Silliman et al., 2019). Thus, rapid restoration of marsh dieback is crucial in order to avoid further degradation, especially in stressed urban salt marshes. The urgency increases when taking into consideration that dieback areas in West Ashley and other studied locations are already associated with lower elevation terrain (Alber et al., 2008; Marsh et al., 2016). Moreover, unvegetated marshes are prone to fill adjacent tidal channels and to experience marsh platform erosion, which is exacerbated during low tide rainfall events (Ganju et al., 2017; Temmerman et al., 2012; Torres et al., 2004). Thus, rapid restoration by grass planting in marshes that have experienced long-term dieback would not only reinstate the marsh’s ability to accrete sediment, but also prevent further disturbance of marsh hydrology. The effectiveness of any restoration effort over the long term must include careful consideration of elevation capital and hydrology. Recent studies show that drainage density and distance from a channel network increase marsh resilience to future drought-induced dieback events (Liu et al., 2020).

### 4.3 Adaptive management of salt marsh ecosystems under climate change

Grass transplanting is an effective restoration practice that bolsters high rates of sediment accretion under conservative SLR scenarios (Silliman et al., 2019; Temmink et al., 2020). However, our model results indicate that even highly productive marshes will likely drown under more extreme climate change scenarios. Our results are supported by long-term modelling performed in Georgia, USA which predicts an 88 % loss of initial marsh area by 2160 (Langston et al., 2021). Field verification of remotely sensed data and site-specific monitoring are critical for the assessment of salt marsh resilience to accelerated SLR in urban environments. If grass planting alone is not sufficient to maintain marsh elevation under accelerated SLR, natural accretion could be augmented by the direct application of sediment. This can be performed by either beneficial use of dredging material, or by sediment redistribution within the same marsh system from edge marsh erosion, or other processes (Ford et al., 1999; Hopkinson et al., 2018). However, even periodic placement of dredged material may not be sufficient under extreme climate change scenarios (Fig 6). Thus, constant monitoring, rapid restoration, and climate change mitigation actions are all urgently needed to preserve highly valuable present-day marsh ecosystems close to urban environments.

## 5. Acknowledgements

This research was funded by grant 66115 (to J.E.K.) from the National Fish and Wildlife Foundation under the National Coastal Resilience Fund program. The views and conclusions contained in this document are those of the authors and should not be interpreted as representing the opinions or policies of the U.S. Government or the National Fish and Wildlife Foundation and its funding sources. Mention of trade names or commercial products does not constitute their endorsement by the U.S. Government, or the National Fish and Wildlife Foundation or its funding sources. Grass planting in Maryville, South Carolina was implemented through funding received from NOAA National Marine Fisheries Service (to M.H.). This research was also funded by grant 1654853 (to J.T.M.) from the National Science Foundation’s Division of Environmental Biology

## 7. Contributions

J.LR., J.E.K., M.H., N.W., J.R., A.G., J.C., and J.T.M. designed the research. J.L.R., M.H., K.D.G., G.K., N.W., J.R., and J.E.K performed field work, J.L.R., and M.H., conducted laboratory work, J.L.R. analyzed the data with input from J.T.M., and J.E.K. J.L.R. wrote the first draft of the manuscript with inputs from J.E.K. and all authors equally contributed to revisions and gave approval for publication

## 8. Data availability statement

Remote sensed imagery was retrieved from publicly available repositories (https://earthexplorer.usgs.gov/ and https://planet.com/). All code and field data are publicly available at a Zenodo repository: https://doi.org/10.5281/zenodo.6980907

## Supplementary Information

### Supplementary Text S1

#### Correction of LIDAR derived DEM

Error correction of the 2017 light detection and ranging (LIDAR) derived digital elevation model (DEM) was constructed based on a multiple regression model (further detail in Materials and Methods). Error was determined based on the difference between real-time kinematic (RTK) survey points and values from the LIDAR derived DEM raster. Inputs for the model selection were NDVI, MSAVI2, and independent bands from a PlanetScope satellite image taken days before the LIDAR acquisition. Model selection determined the following regression model as the best fit:

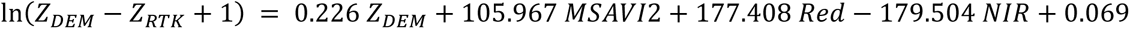

Where Z is elevation in NAVD88 (m).

The adjusted R^2^ in the training dataset was 41.17%

The median (1^st^ – 3^rd^ quartile) error of the original and corrected DEM were -0.03 m (-0.06 – 0.002 m), and -0.01 m (-0.05 – 0.01 m), respectively. Model correction reduced the RMSE of both the test and train datasets in 27.3% and 30.5%, respectively (Fig. S2). The RMSE of the corrected test and train dataset were of 5.7 cm, and 8.0 cm, respectively. These values are withing the range of other error corrected DEMs. For instance, RMSE values from a corrected DEM in Sapelo Island, GA ranged from 5 cm to 30 cm depending on the cover class (Hladik and Alber, 2012).

**Supplementary Table S1:**
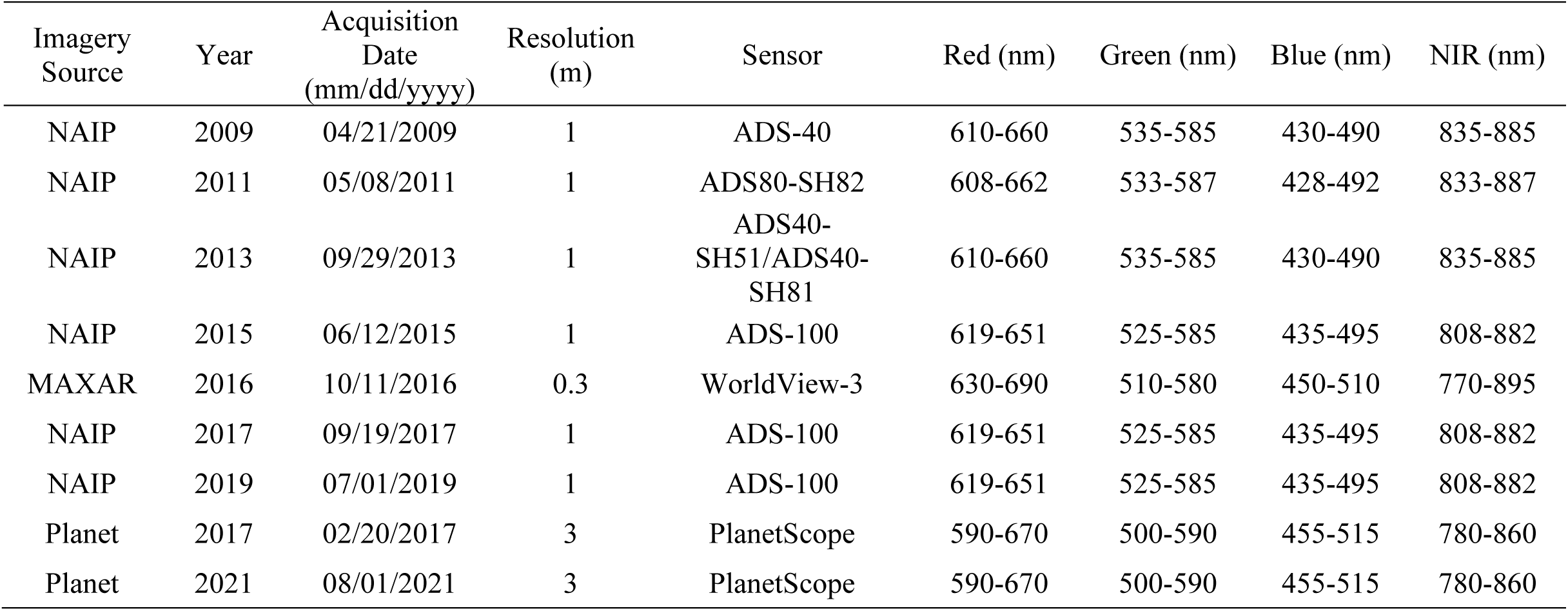
Technical details of aerial and satellite imagery utilized in the present study.

**Supplementary Table S2:**
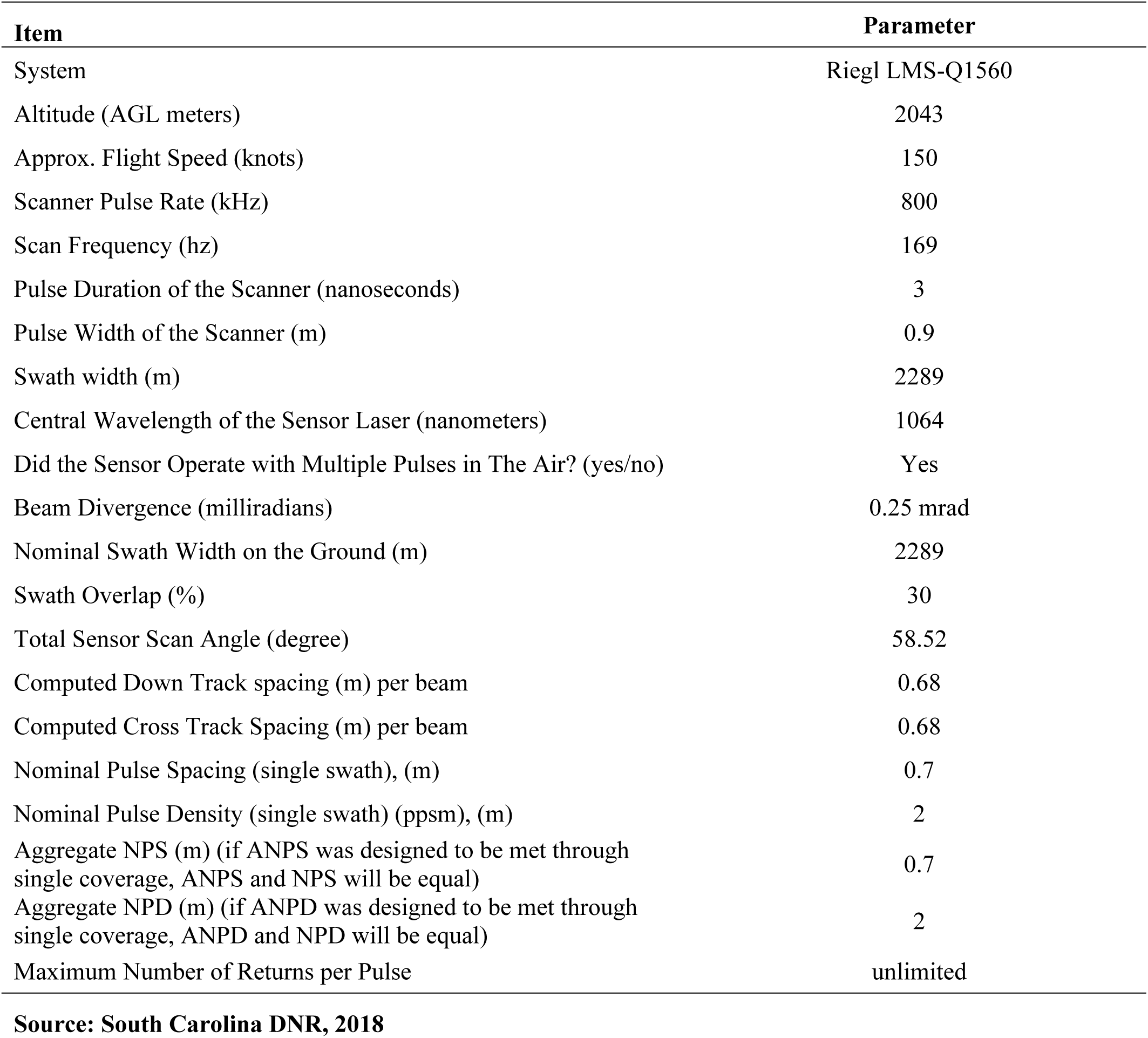
Technical description of the light detection and ranging (LIDAR) acquisition. Information retrieved from South Carolina DNR (2018).

**Supplementary Figure S1.**
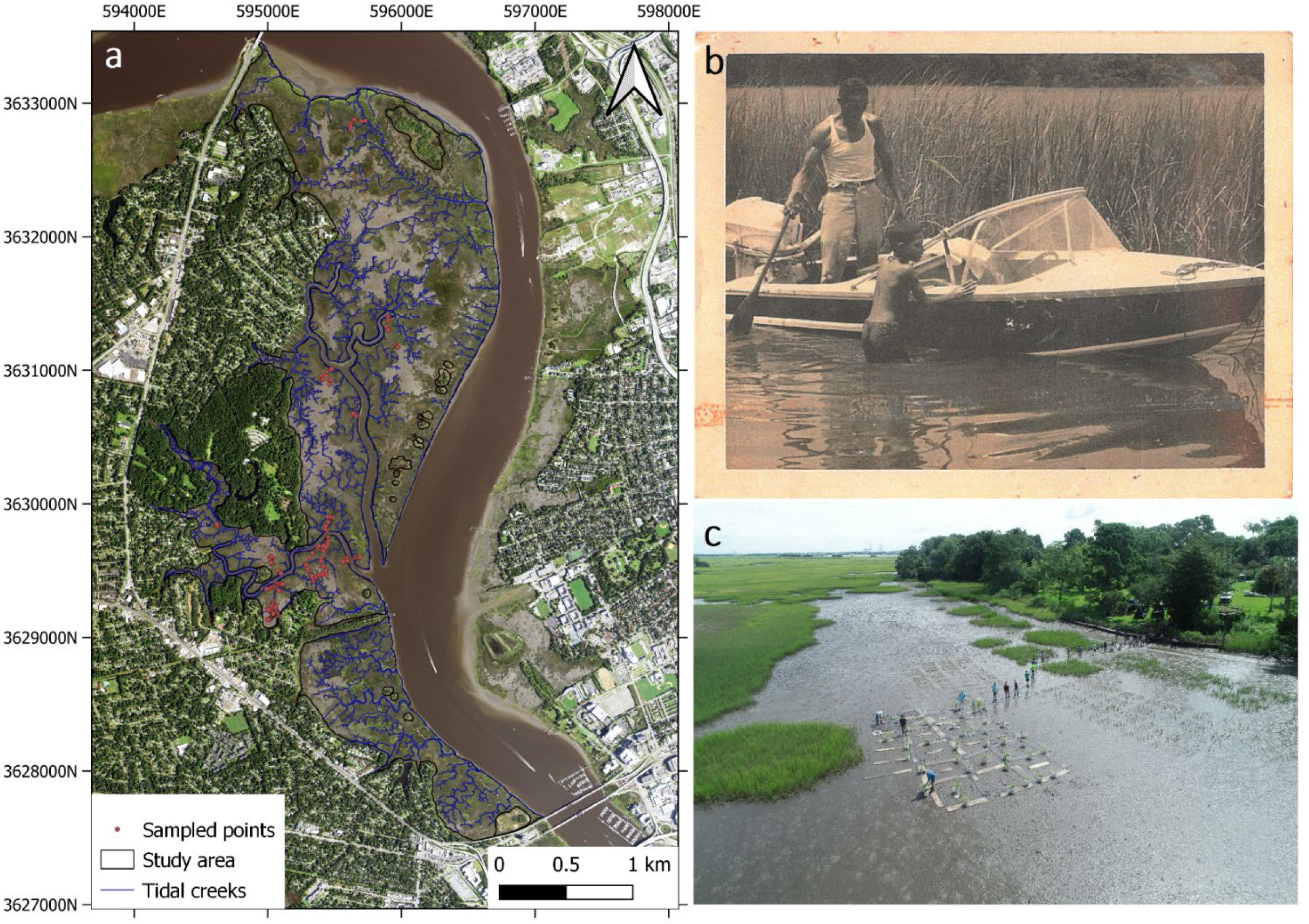
Study area and sampling locations of the present study in West Ashley, Charleston, SC. **a** Map of the study area. Blue lines represent tidal creeks, orange dots sampled location in the field trips of May and July of 2021. **b** Picture of the now dieback Maryville marsh circa 1970. Image provided by Mr. John Carr Jr. **c** Drone image of the Maryville marsh taken on 21 July of 2021. Drone image acquired by the South Carolina Department of Natural Resources Shellfish Research Section.

**Supplementary Figure S2.**
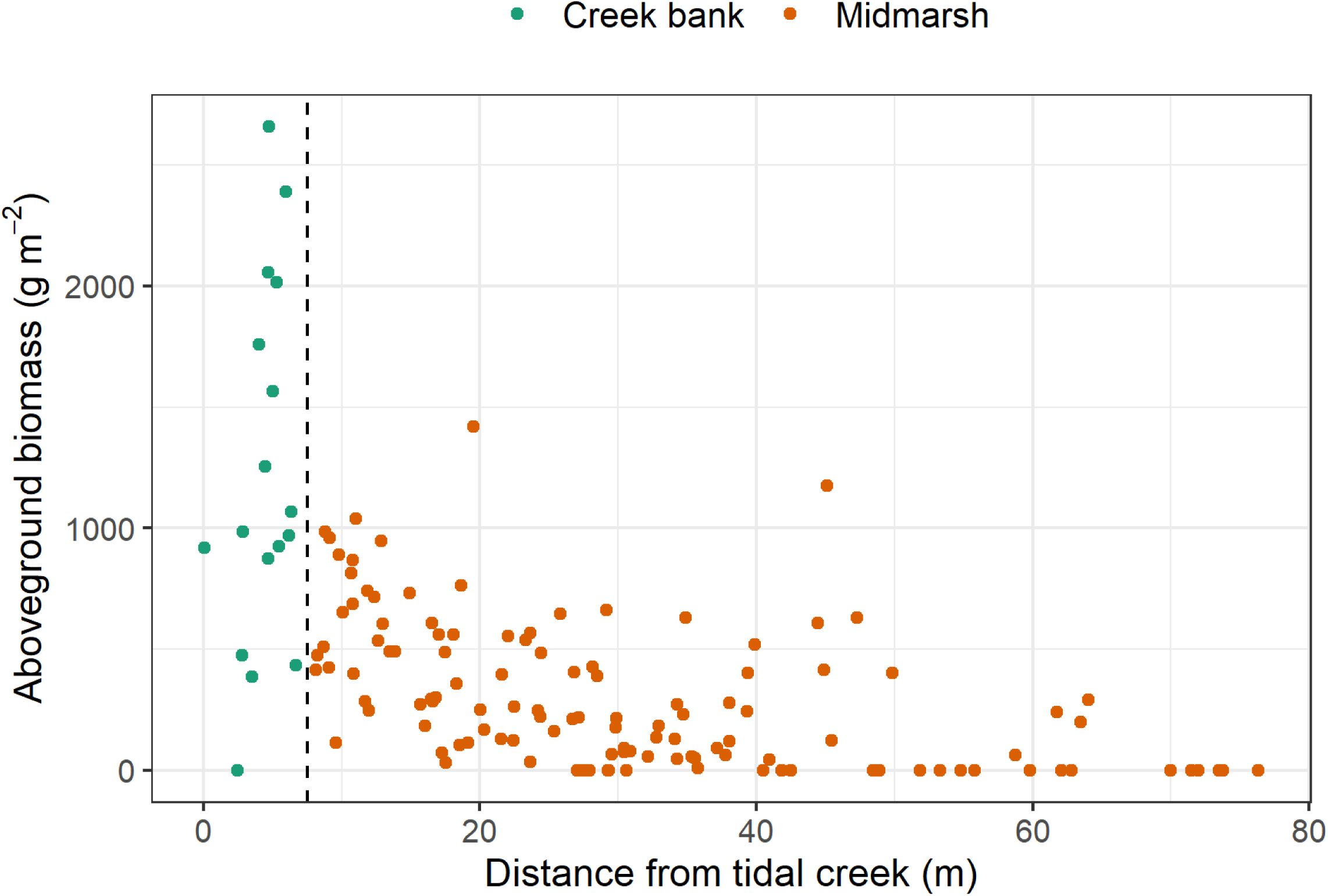
Relationship between *in situ* aboveground biomass and distance from tidal creek. The vertical dotted line represents the tipping point (7.5 m distance from the closest tidal creek) defining the state change between creek bank and midmarsh *Spartina alterniflora* population. Vegetation data was generated during the field trip of July of 2021.

**Supplementary Figure S3:**
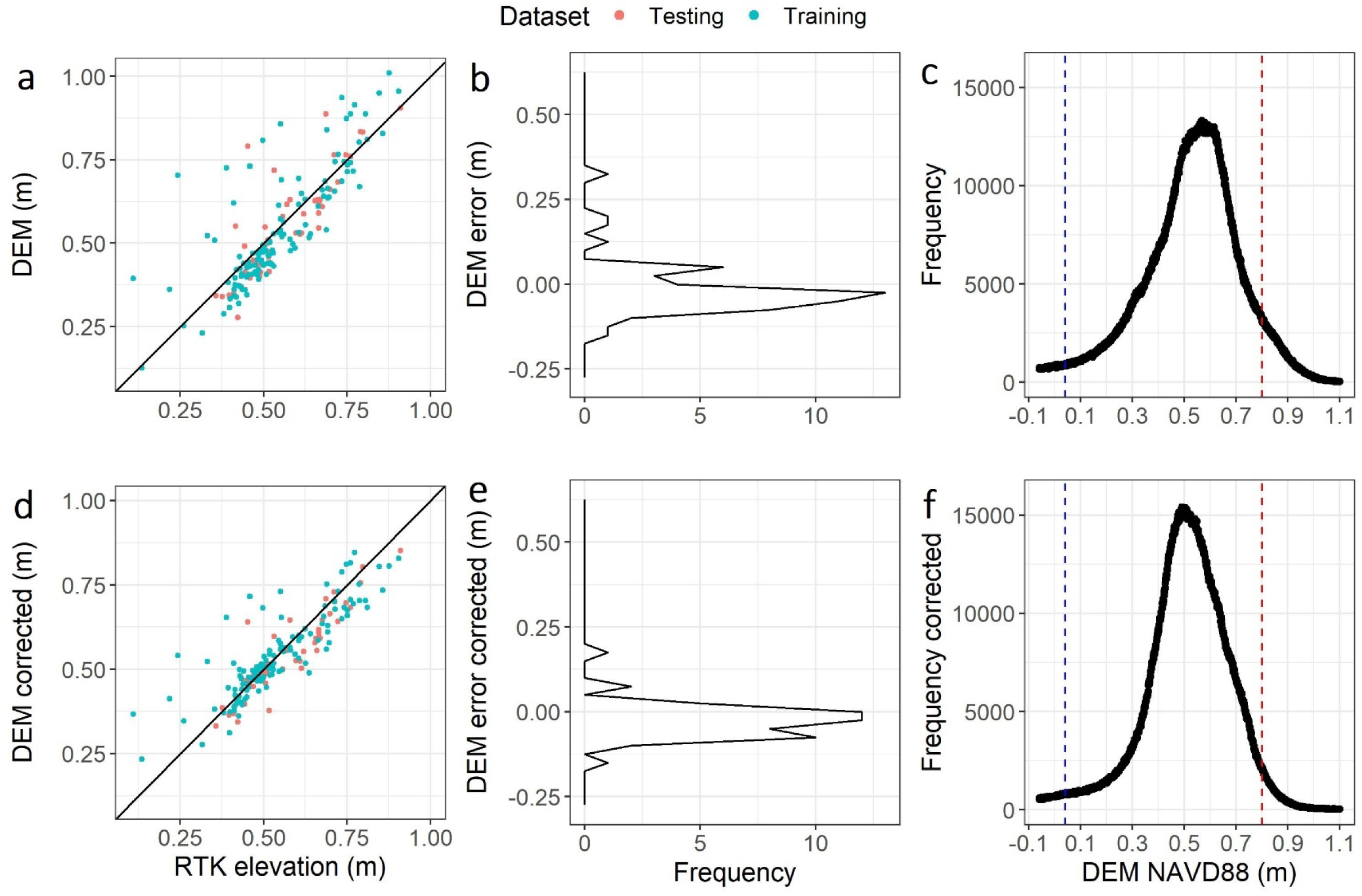
Error correction of the 2017 light detection and ranging (LIDAR) derived digital elevation model (DEM). Biplot of the observed elevation from a real time-kinematic (RTK) survey and elevation from the uncorrected (**a**) and corrected (**d**) 2017 LIDAR derived DEM. Frequency profile of the DEM error (LIDAR elevation – RTK elevation) of the uncorrected (**b**) and corrected (**e**) dataset. Frequency profile of the uncorrected (**c**) and corrected (**f**) elevation from the 2017 LIDAR derived DEM.

**Supplementary Figure S4.**
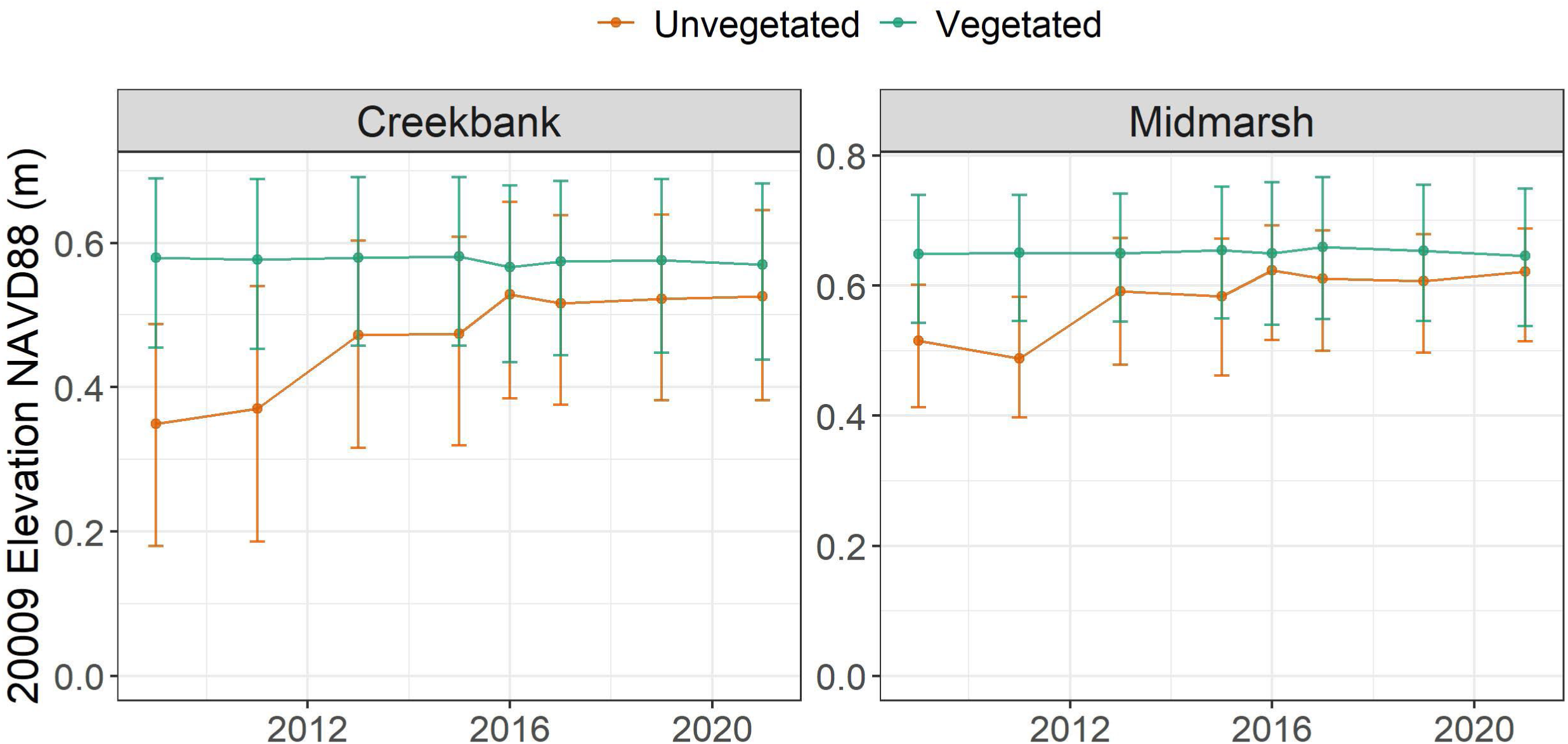
Median ± interquartile range of an uncorrected 2009 light detection and ranging (LIDAR) derived digital elevation model (DEM) of vegetated and unvegetated areas over time (South Carolina DNR, 2009).

