## Supplementary Information for "Restoration and resilience to sea level rise of a salt marsh affected by dieback events in Charleston, SC"

**Supplementary Text S1**

Correction of LIDAR derived DEM.

Error correction of the 2017 light detection and ranging (LIDAR) derived digital elevation model (DEM) was constructed based on a multiple regression model (further detail in Materials and Methods). Error was determined based on the difference between real-time kinematic (RTK) survey points and values from the LIDAR derived DEM raster. Inputs for the model selection were NDVI, MSAVI2, and independent bands from a PlanetScope satellite image taken days before the LIDAR acquisition. Model selection determined the following regression model as the best fit:

$$\ln\left( Z_{DEM}-Z_{RTK}+1 \right) = {0.226 Z}_{DEM}+105.967 MSAVI2+177.408 Red-179.504 NIR+0.069$$

Where Z is elevation in NAVD88 (m).

The adjusted R^2^ in the training dataset was 41.17%

The median (1^st^ – 3^rd^ quartile) error of the original and corrected DEM were -0.03 m (-0.06 – 0.002 m), and -0.01 m (-0.05 – 0.01 m), respectively. Model correction reduced the RMSE of both the test and train datasets in 27.3% and 30.5%, respectively (Fig. S2). The RMSE of the corrected test and train dataset were of 5.7 cm, and 8.0 cm, respectively. These values are withing the range of other error corrected DEMs. For instance, RMSE values from a corrected DEM in Sapelo Island, GA ranged from 5 cm to 30 cm depending on the cover class (Hladik and Alber, 2012).

**Supplementary Table S1:** Technical details of aerial and satellite imagery utilized in the present study.

| Imagery Source | Year | Acquisition Date (mm/dd/yyyy) | Resolution (m) | Sensor | Red (nm) | Green (nm) | Blue (nm) | NIR (nm) |
| --- | --- | --- | --- | --- | --- | --- | --- | --- |
| NAIP | 2009 | 04/21/2009 | 1 | ADS-40 | 610-660 | 535-585 | 430-490 | 835-885 |
| NAIP | 2011 | 05/08/2011 | 1 | ADS80-SH82 | 608-662 | 533-587 | 428-492 | 833-887 |
| NAIP | 2013 | 09/29/2013 | 1 | ADS40-SH51/ADS40-SH81 | 610-660 | 535-585 | 430-490 | 835-885 |
| NAIP | 2015 | 06/12/2015 | 1 | ADS-100 | 619-651 | 525-585 | 435-495 | 808-882 |
| MAXAR | 2016 | 10/11/2016 | 0.3 | WorldView-3 | 630-690 | 510-580 | 450-510 | 770-895 |
| NAIP | 2017 | 09/19/2017 | 1 | ADS-100 | 619-651 | 525-585 | 435-495 | 808-882 |
| NAIP | 2019 | 07/01/2019 | 1 | ADS-100 | 619-651 | 525-585 | 435-495 | 808-882 |
| Planet | 2017 | 02/20/2017 | 3 | PlanetScope | 590-670 | 500-590 | 455-515 | 780-860 |
| Planet | 2021 | 08/01/2021 | 3 | PlanetScope | 590-670 | 500-590 | 455-515 | 780-860 |

**Supplementary Table S2**: Technical description of the light detection and ranging (LIDAR) acquisition. Information retrieved from South Carolina DNR (2018).

| **Item** | **Parameter** |
| --- | --- |
| System | Riegl LMS-Q1560 |
| Altitude (AGL meters) | 2043 |
| Approx. Flight Speed (knots) | 150 |
| Scanner Pulse Rate (kHz) | 800 |
| Scan Frequency (hz) | 169 |
| Pulse Duration of the Scanner (nanoseconds) | 3 |
| Pulse Width of the Scanner (m) | 0.9 |
| Swath width (m) | 2289 |
| Central Wavelength of the Sensor Laser (nanometers) | 1064 |
| Did the Sensor Operate with Multiple Pulses in The Air? (yes/no) | Yes |
| Beam Divergence (milliradians) | 0.25 mrad |
| Nominal Swath Width on the Ground (m) | 2289 |
| Swath Overlap (%) | 30 |
| Total Sensor Scan Angle (degree) | 58.52 |
| Computed Down Track spacing (m) per beam | 0.68 |
| Computed Cross Track Spacing (m) per beam | 0.68 |
| Nominal Pulse Spacing (single swath), (m) | 0.7 |
| Nominal Pulse Density (single swath) (ppsm), (m) | 2 |
| Aggregate NPS (m) (if ANPS was designed to be met through single coverage, ANPS and NPS will be equal) | 0.7 |
| Aggregate NPD (m) (if ANPD was designed to be met through single coverage, ANPD and NPD will be equal) | 2 |
| Maximum Number of Returns per Pulse | unlimited |
| **Source: South Carolina DNR, 2018** |  |

**Supplementary Figure S1.** Study area and sampling locations of the present study in West Ashley, Charleston, SC. **a** Map of the study area. Blue lines represent tidal creeks, orange dots sampled location in the field trips of May and July of 2021. **b** Picture of the now dieback Maryville marsh circa 1970. Image provided by Mr. John Carr Jr. **c** Drone image of the Maryville marsh taken on 21 July of 2021. Drone image acquired by the South Carolina Department of Natural Resources Shellfish Research Section.

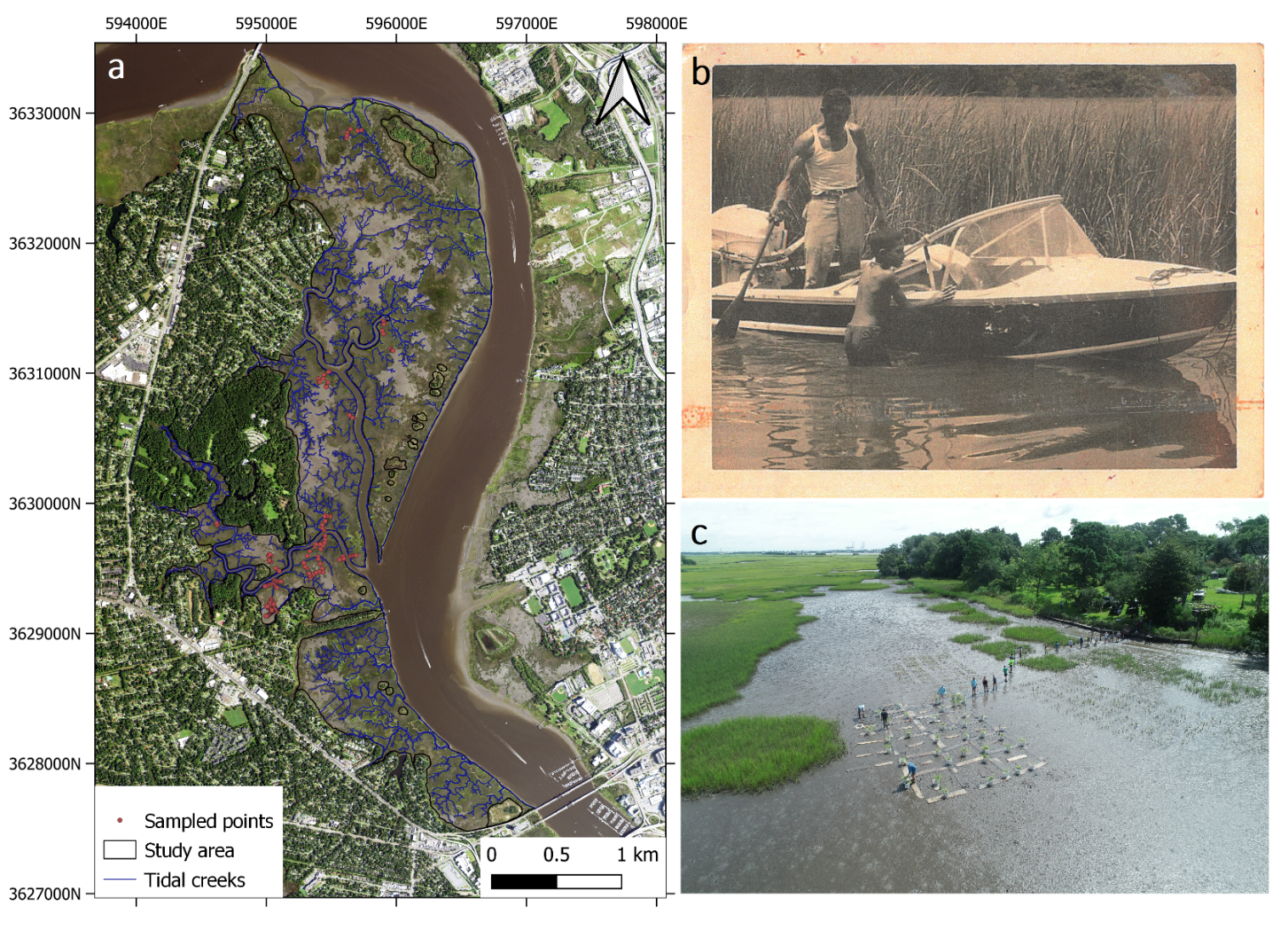

**Supplementary Figure S2.** Relationship between *in situ* aboveground biomass and distance from tidal creek. The vertical dotted line represents the tipping point (7.5 m distance from the closest tidal creek) defining the state change between creek bank and midmarsh *Spartina alterniflora* population. Vegetation data was generated during the field trip of July of 2021.

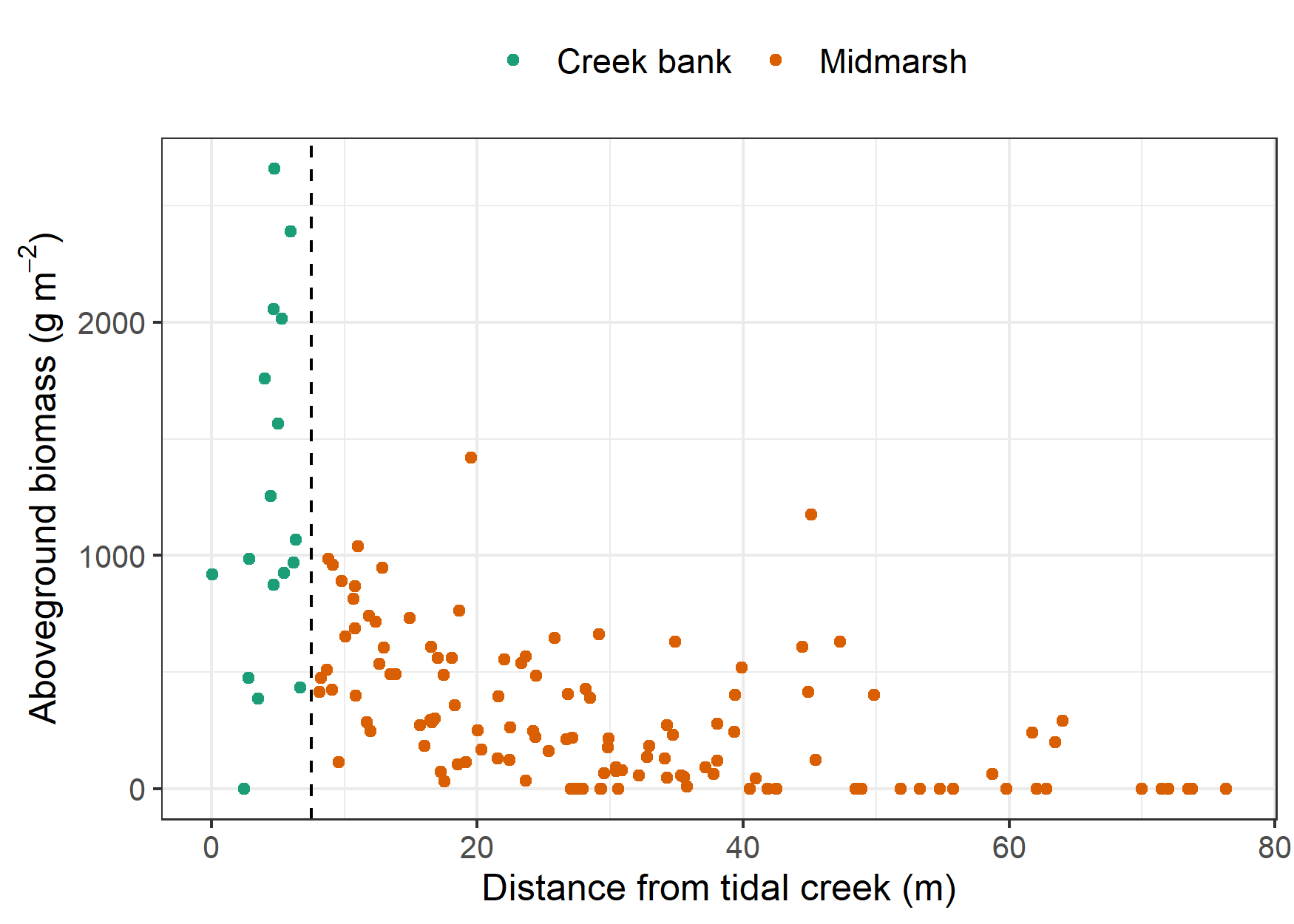

**Supplementary Figure S3:** Error correction of the 2017 light detection and ranging (LIDAR) derived digital elevation model (DEM). Biplot of the observed elevation from a real time-kinematic (RTK) survey and elevation from the uncorrected (**a**) and corrected (**d**) 2017 LIDAR derived DEM. Frequency profile of the DEM error (LIDAR elevation – RTK elevation) of the uncorrected (**b**) and corrected (**e**) dataset. Frequency profile of the uncorrected (**c**) and corrected (**f**) elevation from the 2017 LIDAR derived DEM.

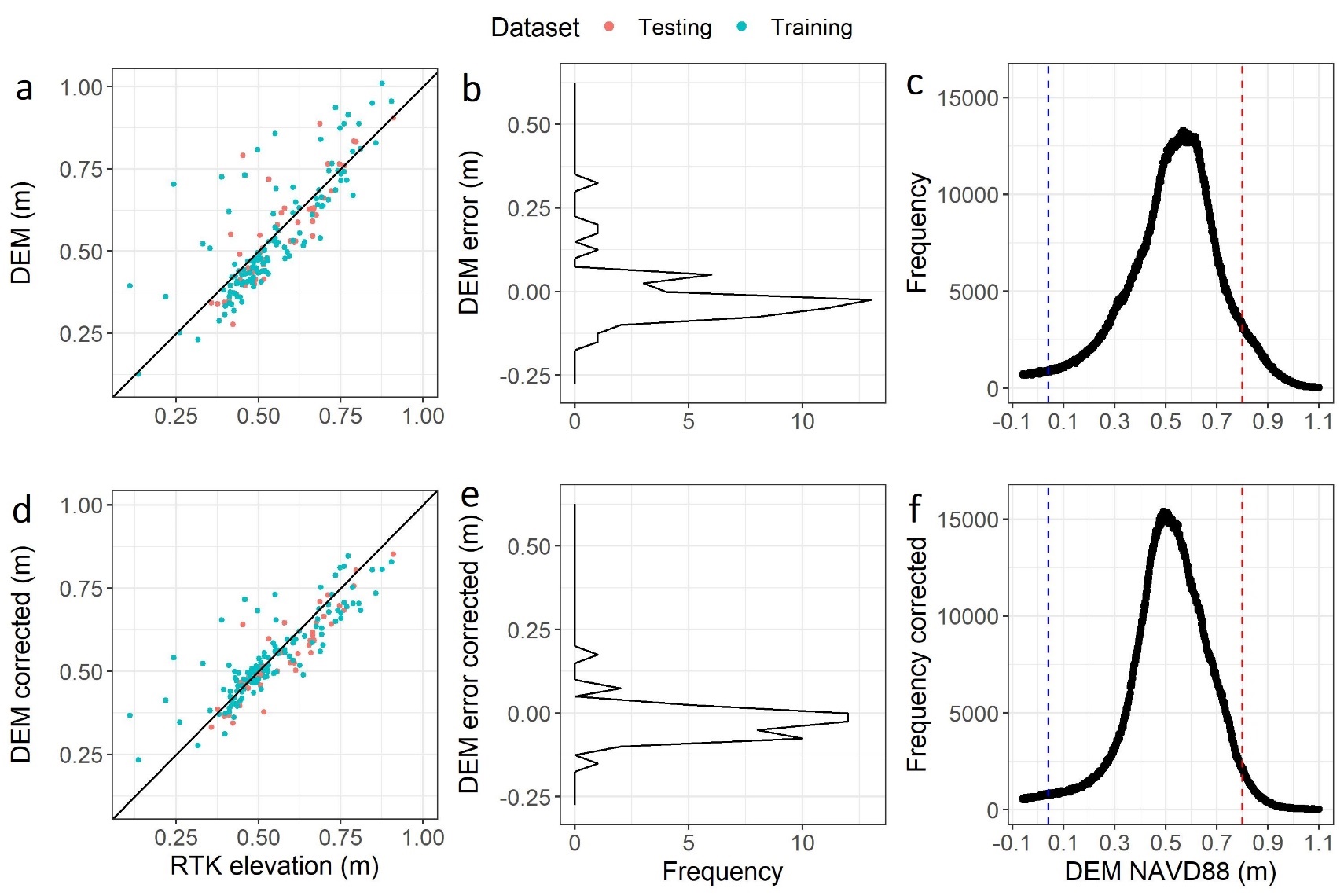

**Supplementary Figure S4.** Median ± interquartile range of an uncorrected 2009 light detection and ranging (LIDAR) derived digital elevation model (DEM) of vegetated and unvegetated areas over time (South Carolina DNR, 2009).

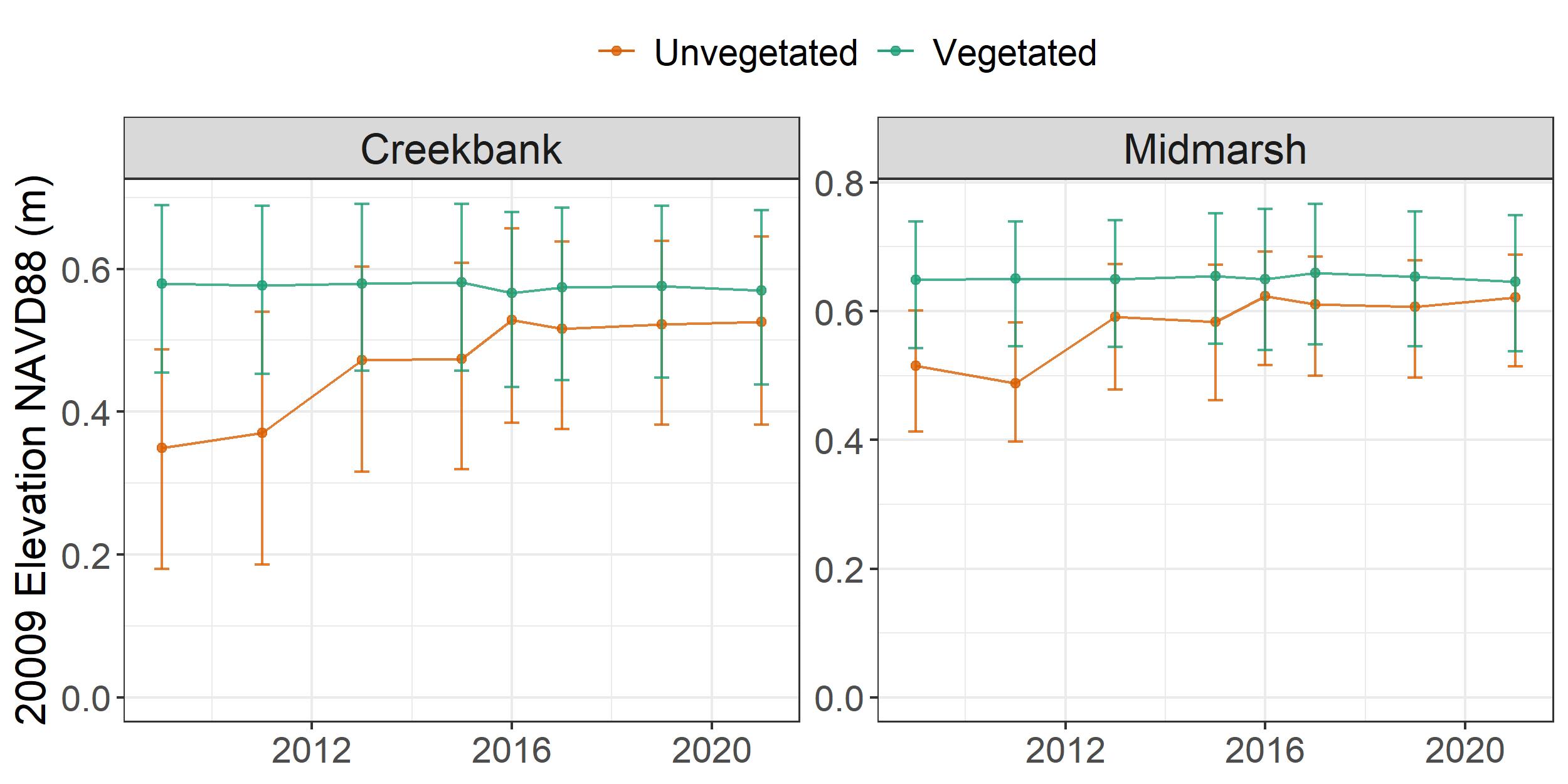
